# A validation framework for neuroimaging software: the case of population receptive fields

**DOI:** 10.1101/2020.01.07.897991

**Authors:** Garikoitz Lerma-Usabiaga, Noah Benson, Jonathan Winawer, Brian A. Wandell

## Abstract

Neuroimaging software methods are complex, making it a near certainty that some implementations will contain errors. Modern computational techniques (i.e., public code and data repositories, continuous integration, containerization) enable the reproducibility of the analyses and reduce coding errors, but they do not guarantee the scientific validity of the results. It is difficult, nay impossible, for researchers to check the accuracy of software by reading the source code; ground truth test datasets are needed. Computational reproducibility means providing software so that for the same input anyone obtains the same result, right or wrong. Computational validity means obtaining the right result for the ground-truth test data. We describe a framework for validating and sharing software implementations. We apply the framework to an application: population receptive field (pRF) methods for functional MRI data. The framework is composed of three main components implemented with containerization methods to guarantee computational reproducibility: (1) synthesis of fMRI time series from ground-truth pRF parameters, (2) implementation of four public pRF analysis tools and standardization of inputs and outputs, and (3) report creation to compare the results with the ground truth parameters. We identified realistic conditions that lead to imperfect parameter recovery in all four implementations, and we provide means to reduce this problem. The computational validity framework supports scientific rigor and creativity, as opposed to the oft-repeated suggestion that investigators rely upon a few agreed upon packages. The framework and methods can be extended to other critical neuroimaging algorithms. Having validation frameworks help (1) developers to build new software, (2) research scientists to verify the software’s accuracy, and (3) reviewers to evaluate the methods used in publications and grants.

**Author Summary:** Computer science provides powerful tools and techniques for implementing and deploying software. These techniques support software collaboration, reduce coding errors and enable reproducibility of the analyses. A further question is whether the software estimates are correct (valid). We describe a framework for validating and sharing software implementations based on ground-truth testing. We applied the framework to four separate applications that implemented population receptive field (pRF) estimates for functional MRI data. We quantified the validity, and we also documented limitations with these applications. Finally, we provide ways to mitigate these limitations. Implementing a software validation framework along with sharing and reproducibility is an important step for the complex methods used in neuroscience. Validation will help developers to build new software, researchers verify that the results are valid, and reviewers to evaluate the precision of methods in publications and grants.

## Introduction

Neuroimaging software methods are based on complex architectures, thousands of lines of code, and hundreds of configuration parameters. Consequently, it is a near certainty that some implementations will contain errors. Modern computational techniques (i.e. public code and data repositories, continuous integration, containerization) enable the reproducibility of the analyses and the reduction of coding errors, but do not guarantee the scientific validity of the results.

Computational reproducibility—enabling anyone to obtain the same result for the same input data and code—is one important component of scientific reproducibility [1,2]. Computational generalization means obtaining the same result for the same input data and different software implementations. Computational validity is the further test as to whether the result is correct. For reviewers and scientists alike, it is impossible to establish validity by just reading code. Scientific journal readers assume that the tool provides correct results if it was correctly set-up and used, but this is not guaranteed. To establish validity, ground truth test datasets are required. In this way we can establish that the results are valid over a well-defined range of inputs (scope).

Here, we describe a computational framework for validating and sharing software implementations (Figure 1). The framework is divided into three parts. The x-Synthesize part comprises methods to produce synthetic test data with known parameters in a defined file format. The x-Analysis part uses the test data as inputs to the algorithms under test. These algorithms are incorporated in containers that accept test data inputs and produce output files in a well-defined format. The x-Reporting tools compare the outputs with the ground-truth parameters in the x-Synthesize part. The x-Analysis containers can also analyze experimental rather than synthetic data placed in the input file format. This framework permits the user to compare multiple algorithms in different containers.

**Figure 1.**
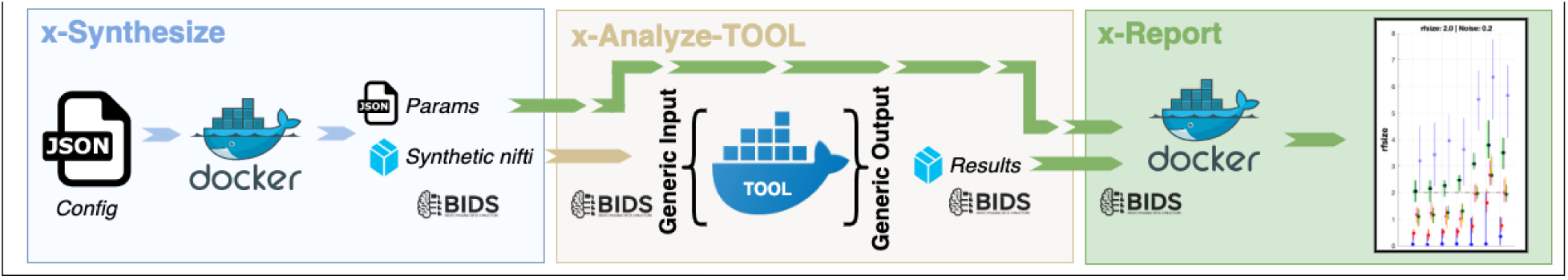
Validation framework overview. The framework is composed of three main components. *x-Synthesize:* synthesis of neuroimaging data from ground truth parameters; *x-Analyze-TOOL:* implementation of analysis tools and standardization of inputs and outputs; *x-Report* compares the tool outputs the ground truth parameters. The three components are implemented within containers. Container parameters are specified by JSON text files.

We use ‘x’ as a placeholder that is replaced by the name of a specific analysis. The framework and methods can be extended to many neuroimaging tools. In this paper, we implemented population receptive field (pRF) methods for functional MRI data. The main motivation of beginning with pRF methods is the possibility of accurately generating synthetic data because the pRF is a model. Hence, we can ask how accurately the software recovers the model parameters. Following the guidelines, the pRF validity framework is composed of three main components: (1) synthesis of fMRI time series from ground truth pRF parameters, (2) implementation of four pRF analysis tools and standardization of inputs and outputs, and (3) tool-independent report creation to compare the results with the ground truth parameters. The different components are implemented using containerization methods to guarantee computational reproducibility. All components and testing datasets are publicly available. A user can run the framework with only Docker and a text editor installed.

### Related literature

Validating neuroimaging software has been approached in several different ways. We can consider MRI phantoms as the first validation system. Phantoms, commonly used in quantitative MRI software development, provide ground truth data, [3–5].

There have been other efforts in diffusion weighted imaging (DWI) set up in the form of public challenges, such as the Tractometer [6–8], an online tract validation system based on the HCP dataset [9] and a simulated DWI diffusion generated with the tool Fiberbox (http://docs.mitk.org/2014.10/org_mitk_views_fiberfoxview.html). There have been efforts to synthesize fMRI data, some for generic task activations [10–12], and some others more specifically targeted, for example in studies of scotomas [13,14] or temporal integration [15]. See here for a review of functional MRI simulations [16].

## Methods

### Architecture

The software we developed comprises two main components: the open-source code repository (https://github.com/vistalab/prfmodel) and the containers (Docker/Singularity). The containers we implemented for the population receptive field analysis can be installed with one command (‘docker pull’). Users who prefer Singularity can convert the containers from Docker to Singularity (https://github.com/singularityhub/docker2singularity). The input/output of every element of the framework follows the Brain Imaging Data Structure (BIDS) format [17]. This means that x-Synthesize will generate synthetic data in BIDS format, and that the output of the x-Analyze and x-Report will be in BIDS derivatives format.

The following sections describe the three parts of the framework. The case we describe tests population receptive field (pRF) algorithms: mrVista (https://github.com/vistalab/vistasoft), AFNI (https://github.com/afni/afni), Popeye (https://github.com/kdesimone/popeye), analyzePRF (https://github.com/kendrickkay/analyzePRF). These four are a subset of the public implementations (https://github.com/vistalab/PRFmodel/wiki/Public-pRF-implementations). It is our intention for the framework to be applicable to many other analyses.

### Implementation

#### Synthesize

The prf-Synthesize container generates synthetic BOLD time series data. We use the naming convention x-Synthesize where ‘x’ is replaced by the algorithm under test. For example, glm-Synthesize, dti-Synthesize, csd-Synthesize, and so forth. See the Appendix for a basic *How to use* guide.

The synthetic BOLD signal is created using the forward model that is implicit when searching for pRF parameters. The forward model is comprised of several components (Figure 2). The stimulus is represented by a sequence of 2D binary images that represent the presence (1) or absence (0) of contrast at each location. The two-dimensional receptive field (RF) is also represented as a matrix; the inner product of the stimulus and the RF matrix produces a response time series. The time series is convolved with the hemodynamic response function (HRF) to obtain the noise-free BOLD signal. A noise model produces a signal that is added, generating the noise-free BOLD signal.

**Figure 2.**
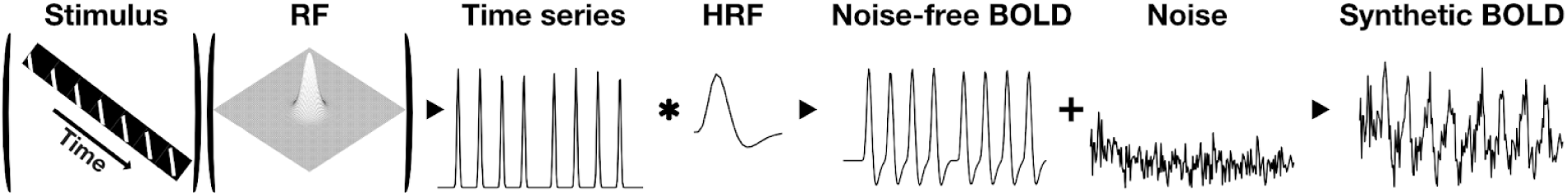
prf-Synthesize: Simulation of ground-truth BOLD data with a forward model. The prf-Synthesize container reads the JSON parameter file to define the RF parameters, and a NIfTI file to specify the stimulus contrast time series. The noise-free BOLD signal is calculated from the inner product of the stimulus and RF, which generates a time series. This time series is convolved with the HRF to obtain the noise-free BOLD signal. A parameterized noise model produces a signal that is added to generate the synthetic BOLD signal. *RF: Receptive field; HRF: Hemodynamic response function. BOLD: blood-oxygen-level dependent*

Each of these components is controlled by explicit parameters that are available through the prf-Synthesize interface. The outputs at each stage of the calculation can be visualized and modified if the user chooses to install the Matlab code (see Appendix for details).

#### Receptive Field

The population receptive field is represented as a 2D function over the input image (stimulus). The simplest and most widely used pRF models are implemented in prf-Synthesize (Figure 2-RF). The Gaussian receptive field has five parameters, :*G*_*i*_(*x, y, σ*_1_, *σ*_2_, *θ*):

- The center position of the receptive field (x, y).
- The standard deviations (*σ*_*i*_) of the two axes of the ellipse
- The angle (*θ*) of the main axis (larger sigma). (Zero for a circular receptive field, *σ*_1_ = *σ*_2_.)

In addition to the Gaussian, we implemented a Difference of Gaussians (DoG) model. In this case, there is also a relative amplitude for the center and surround Gaussians

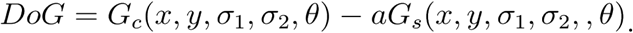

#### Stimulus

We generate the stimulus using the methods described on this web page (https://kendrickkay.net/analyzePRF/). By default, we create a binary stimulus that represents the locations where there is non-zero stimulus contrast. The default contrast map represents 20 deg of visual angle and is stored as a (row, col, time) tensor with 101 row and column samples and 200 temporal samples. These parameters can be varied by adjusting the parameters of prf-Synthesize, specified in the json file. The default stimulus is a bar that sweeps through the visual field in multiple directions and includes some blank periods. The default timing parameters are set to a 200 sec presentation with a sampling of 1 sec.

#### Hemodynamic Response Function (HRF)

The default HRF is the sum of two Gamma probability distribution functions, *x*^(*a*−1)^ exp(−*βx*). Each Gamma distribution has two parameters, and a third parameter defines the relative scaling. As we show below, it is quite significant that different tools implement different HRFs, either in their parameters or in their functional form. We explore the impact of the HRF extensively below.

#### Noise models and parameter selection strategies

We implemented three types of additive noise: white noise, physiological noise (cardiac and respiratory) and low frequency drift noise [10,11,16,18,19]. The level of the white noise is controlled by a single parameter that sets the amplitude. Respiratory, cardiac and low-frequency noises are each controlled by four parameters: frequency, amplitude and a jitter value for frequency and amplitude. The jitter value modifies the frequency and amplitude so that each time the noise model is run the parameters values are randomized.

For the analyses below we simulated the BOLD time series at four noise levels (Figure 3). We selected the noise levels by analyzing the measured BOLD time series from a participant who viewed 8 repetitions of a large-field on/off contrast stimulus. The BOLD data were sampled every 1.5 seconds, each cycle comprised 24 seconds (0.042 Hz) and consisted of 22 axial slices covering the occipital lobe, with 2.5 mm isotropic voxels. The experiment was repeated four times. Three gray matter voxels were selected and the range of noise levels in these voxels were used to set the range of noise levels in the simulation (the fMRI data are available at https://osf.io/3d8rp/).

**Figure 3.**
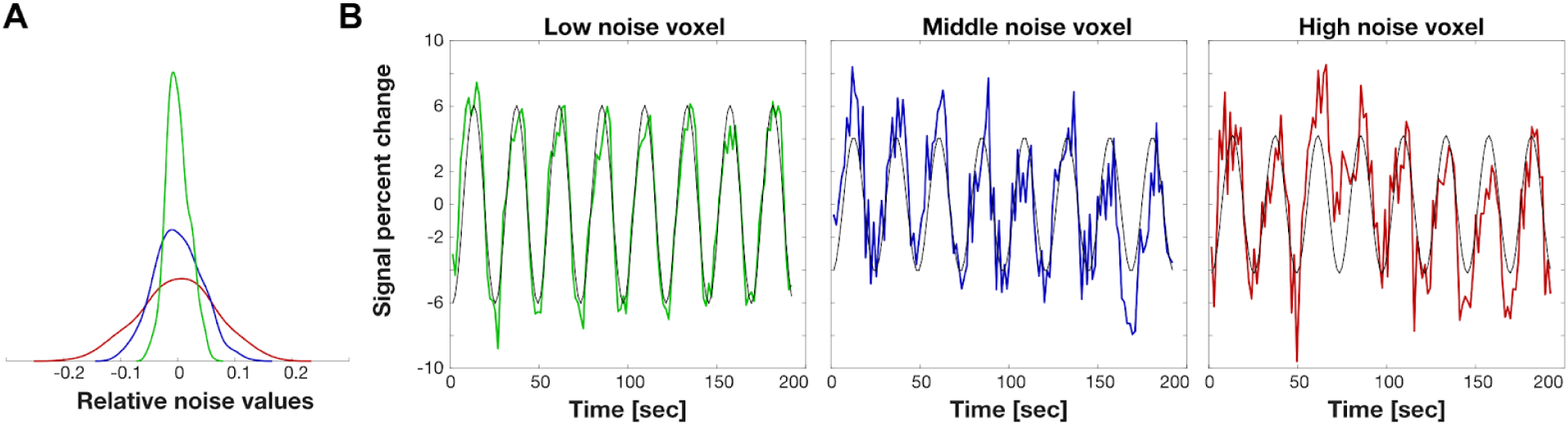
Noise parameter estimation. We analyzed BOLD data collected with an on-off visual contrast stimulus (8 cycles/run, four repetitions). Noise was assessed by comparing the BOLD time series with a harmonic at the same frequency. A) The probability density functions of the noise values of three voxels with low (green), mid (blue) and high (red) noise. B) The colored curves are the mean BOLD time-series plotted as percent modulations (mean of the four repetitions). The black curves are the eight cycle harmonic fit to the data. The noise is the difference between the two. The blue curve is the middle of the noise level found in visual cortex. The other two curves show low and high noise examples that are often found in real measurements. *Low noise: SNR=6*.*55dB; mid noise: SNR=-3*.*03dB; high noise: SNR=-6*.*68dB*.

These three reference voxels were selected using an automated procedure. First, grey matter voxels were identified whose mean BOLD signal exceeded 75% of the total mean BOLD signal. Second, from this set we selected voxels with high local coherence values. The coherence at each voxel was calculated by dividing the amplitude at the stimulus frequency by the mean amplitude of the frequencies in a window around the stimulus frequency. The amplitude spectrum of the BOLD signal of a voxel with high coherence has a local peak at the stimulus frequency, 0.042 Hz. The amplitude at the stimulus frequency in this example is about 10x higher than the expected noise amplitude. The simple on/off stimulus generates significant responses in large regions of the occipital lobe. We selected voxels that responded significantly to the stimulus, at least 80% of the maximum coherence.

Third, the BOLD signal was converted to signal percent change by subtracting and dividing each time series by its mean. We selected the voxels with signal percent change between 8% and 12%. The contrast was calculated as the mean between the maximum and minimum values of the time series. Fourth, noise was measured in the surviving voxels as the difference between the BOLD signal and an 8 cycle sinusoidal fitted to the signal. We selected 3 voxels based on their noise distributions, at 95%, 45% and 10% (Figure 3A) of the voxel with the smallest noise level. Figure 3B shows the mean value of the 3 selected voxel time series and the corresponding fitted sinusoidal. Signal to noise ratio (SNR) was calculated by the ratio of the root mean squared error of the signal (the fitted sinusoidal) and the noise (time series minus the fitted sinusoidal).

Based on these values, we found three parameter settings that correspond to the low- mid - and high-noise levels in the prf-Synthesize tool (Figure 4). These parameters can be set by the user as well. The levels shown in the figure correspond to a single acquisition of the time series. It is common practice to average several scans to increase SNR. For example, averaging 3 mid-noise level time series increases the SNR 2-3 dB.

**Figure 4.**
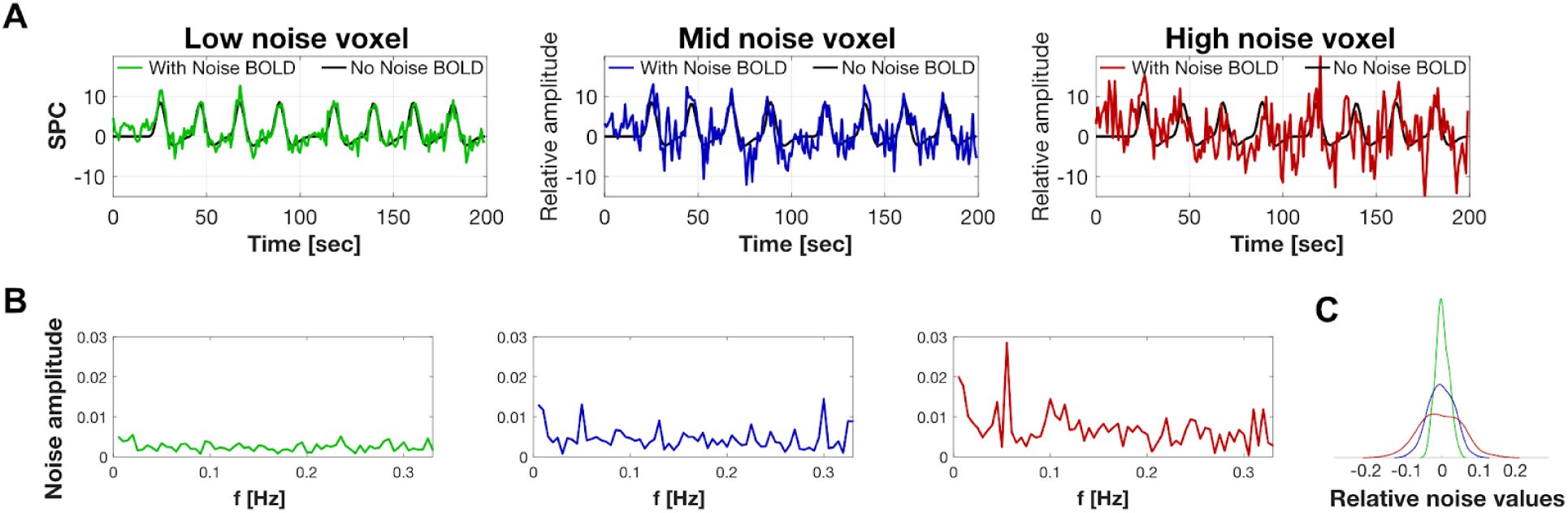
prf-Synthesize BOLD signals calculated for a moving bar retinotopy stimulus. (A) The model calculation of a BOLD signal to a typical moving bar stimulus under the three noise levels. (B) Noise amplitude spectrum for each of the cases in A. (C) The probability density function of the noise values. The simulations match the noise distributions in Figure 3A. *SPC: signal percent change; f: frequency. Low noise: SNR=5*.*29dB; mid noise: SNR=-0*.*51dB; high noise: SNR=-4*.*29dB*

#### Analyze

The prf-Analyze Docker containers implement the pRF tools. The inputs to the container are defined using standard file formats and directory organization. The software repository includes a base Docker container specification so that a pRF tool developer can insert their code and test the implementation using our test framework. To this point, we have implemented four different prf-Analyze containers:

1. **prf-Analyze-vista:** the vistasoft’s pRF Matlab implementation, heavily based on a graphical user interface.
2. **prf-Analyze-afni:** AFNI’s command line based C++ pRF implementation.
3. **prf-Analyze-aprf:** analyzePRF is Kendrick Kay’s PRF Matlab implementation.
4. **prf-Analyze-pop:** Popeye is a pRF python implementation.

Each container takes the stimulus file and the BOLD time series represented as NIfTI files and organized using the BIDS file naming convention and directory tree [17]. This is the format that is generated by prf-Synthesize but could just as well be data from an empirical study. The prf-Synthesize tools are designed to work with BOLD data following general pre-processing, including: removal of the initial volumes to allow longitudinal magnetization to reach steady-state; correction of differences in slice acquisition times; correction for spatial distortion; motion correction. See the Appendix for more information about the prf-Validation User’s Guide.

In the course of implementing and validating the containers, we identified a few issues that needed to be corrected with the original downloads. These issues were reported to the developers and they have updated the code. We note only the issues that might impact results run using the versions in use prior to summer 2019. Three main issues were related to specification of the temporal sampling of the HRF (https://github.com/kdesimone/popeye/pull/64) and the normalization of input BOLD signals (https://github.com/kdesimone/popeye/pull/62) in popeye, and an error in the formula of the ellipse in AFNI (https://afni.nimh.nih.gov/pub/dist/doc/misc/history/afni_hist_BUG_FIX.html). An extension was added to mrVista to enable using synthetic data comprised of one-dimensional NIfTI data format.

#### Report

The third component of the framework, prf-Report container, compares the prf-Analyze-tool outputs with the parameters that specified the ground-truth values used by prf-Synthesize. The input to prf-Report report is a json file that specifies the location of the ground-truth and analyzed data files. The prf-Report container reads these files and creates a summary file that includes the key ground-truth parameters and the corresponding prf-Analyze estimates. The container also generates summary plots that assess algorithm performance. These plots are stored in scalable vector graphics (SVG) format and images in portable network graphics (PNG) format that are readily usable in any report or web-site. See the Appendix for the prf-Validation User’s Guide.

## Results

We analyzed the different tools using synthetic data with and without noise. The main conclusions are observable in the noiseless analysis and also valid for the noise case. The main value of the noise analysis is to illustrate the change in the size of the confidence intervals of the parameter estimates as noise increases.

### Noiseless analysis

#### Accuracy

As a minimum requirement, we expect that the software will return an accurate estimate of the parameters when the synthetic signal is (a) noise-free, and (b) generated using the same assumptions by the analysis, including a matched HRF. Figure 5 compares the four estimates of a circular population RF located at (3,3) deg and with a size of 1 deg.

**Figure 5.**
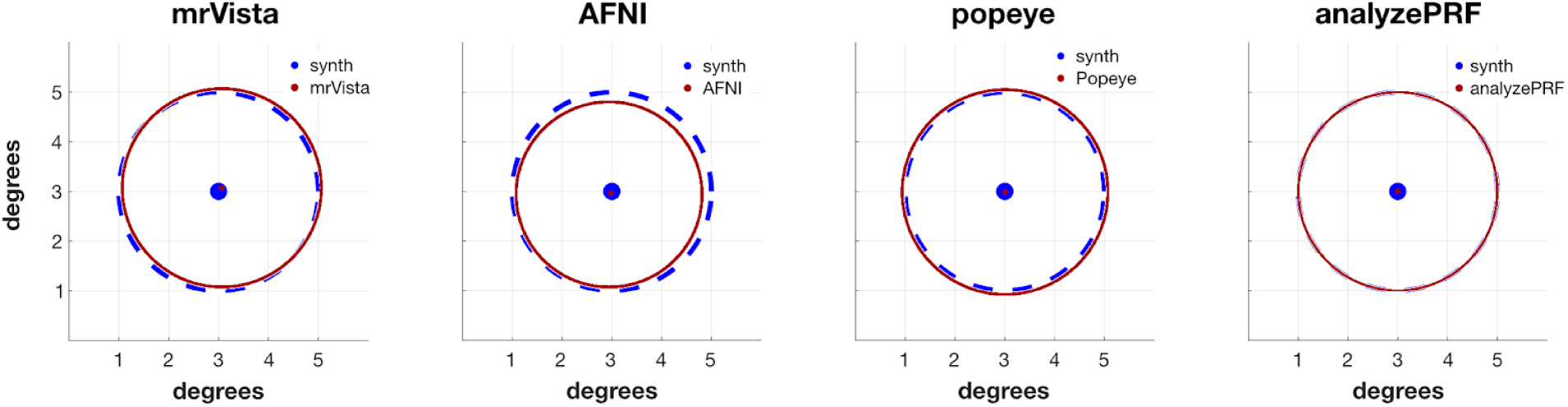
prf-Report for noise-free position and size estimation. prf-Synthesize created BOLD signals from a moving bar stimulus and a gaussian RF centered in x=3 deg, y=3 deg, with a 1 SD radius of 2 deg. Four prf-Synthesize tools estimated the center and size. In each case the noise-free BOLD signals were calculated using the same HRF implemented in the tool. All tools estimated parameters using the linear pRF model.

When downloaded from their sites, the default algorithms differ. For example, some of the algorithms both fit the BOLD data and search to optimize the HRF function. Others include a compressive spatial summation by default. Some of the algorithms permit fitting of more complex pRF shapes (non-circular, difference of Gaussians). Before exploring these additional features - and they should be explored - the validation framework must test the fundamental algorithm of a linear model with a known HRF, which is a common basis for all of the algorithms.

The comparisons in Figure 5 were performed with a base case that all of the tools can analyze: a circular Gaussian pRF modelling, a linear model, and no HRF search [20]. This is a simple test, but we emphasize that this was not generally the default settings when the software was downloaded.

#### HRF mismatch

Every prf-Analyze tool assumes a default HRF. The analysis in Figure 5 was constructed so that the HRF in the prf-Synthesize and prf-Analyze tool matched. For that analysis, we generated a different synthetic BOLD signal for each tool. In this section, we study the performance of the tools when the HRF assumption does not match the prf-Synthesize assumption. We generated BOLD time series using four different HRFs that systematically differed in their width (Figure 6A). We used prf-Validation for these different synthetic data. There is a large and systematic bias in the size estimate that depends on the width of the HRF (Figure 6B). There is a correspondence between the width of the HRF used to synthesize the BOLD signal and the size of calculated pRF: if the width of the HRF used to synthesize is smaller than the width of the HRF assumed by the tool, then the tool will underestimate the size of the pRF (and the opposite is true as well). The Boynton HRF [21] was selected for illustration purposes, because the lack of undershoot made the example clearer, but similar effects were observed with all HRF shapes. We observed mismatch in eccentricity and polar angle as well, but at magnitudes of about a tenth of a degree. For this stimulus and the noise-free case, we consider the HRF effect on polar angle and eccentricity to be negligible.

**Figure 6.**
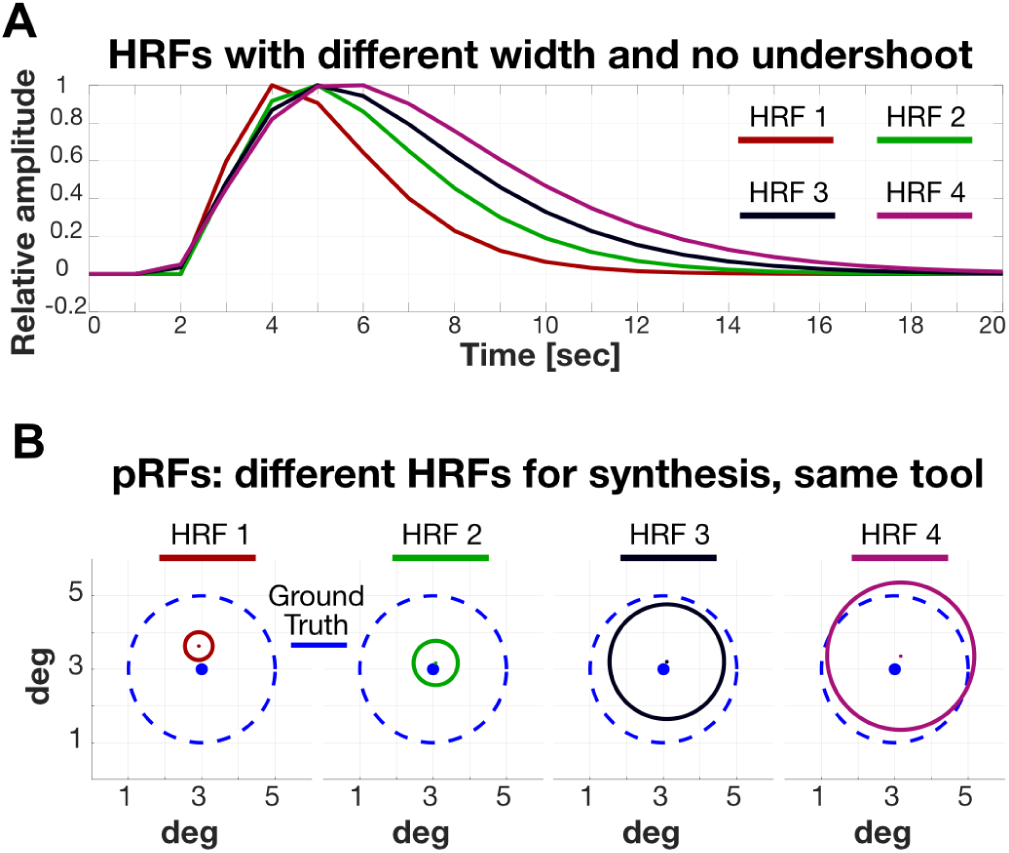
RF size dependence on the HRF shape. (A) Four all-positive HRFs with different widths. (B) The ground-truth signals encoded a circular pRF centered at (3, 3) deg with a radius (1 SD of the Gaussian) of 2 deg (dashed, blue circles). The solid colored circles show the estimated pRF calculated with a representative analysis tool (analyzePRF); all prf-Analyze tools have the same pattern. There is a systematic relationship between the HRF width of the estimated pRF radius.

### Noise analysis

We added three levels (low, mid, high) of white, physiological and low drift noise to the original noise-free signals, and we synthesized 100 repetitions per each. Each repetition was different due to the random nature of the white noise and the jitter introduced to the amplitudes and frequencies. In Figure 7, we show the middle noise case, and illustrate the effect of noise on the pRF estimates (jitter within each plot) and the effect of the relation between the HRF assumed in analysis and the HRF used in the synthesis (differences between plots). We show only the 90% confidence interval of the values, due to the presence of outliers in some of the analysis tools. The centered dash-dot ellipse represents the spread of the centers. The dashed blue curve is ground truth.

**Figure 7.**
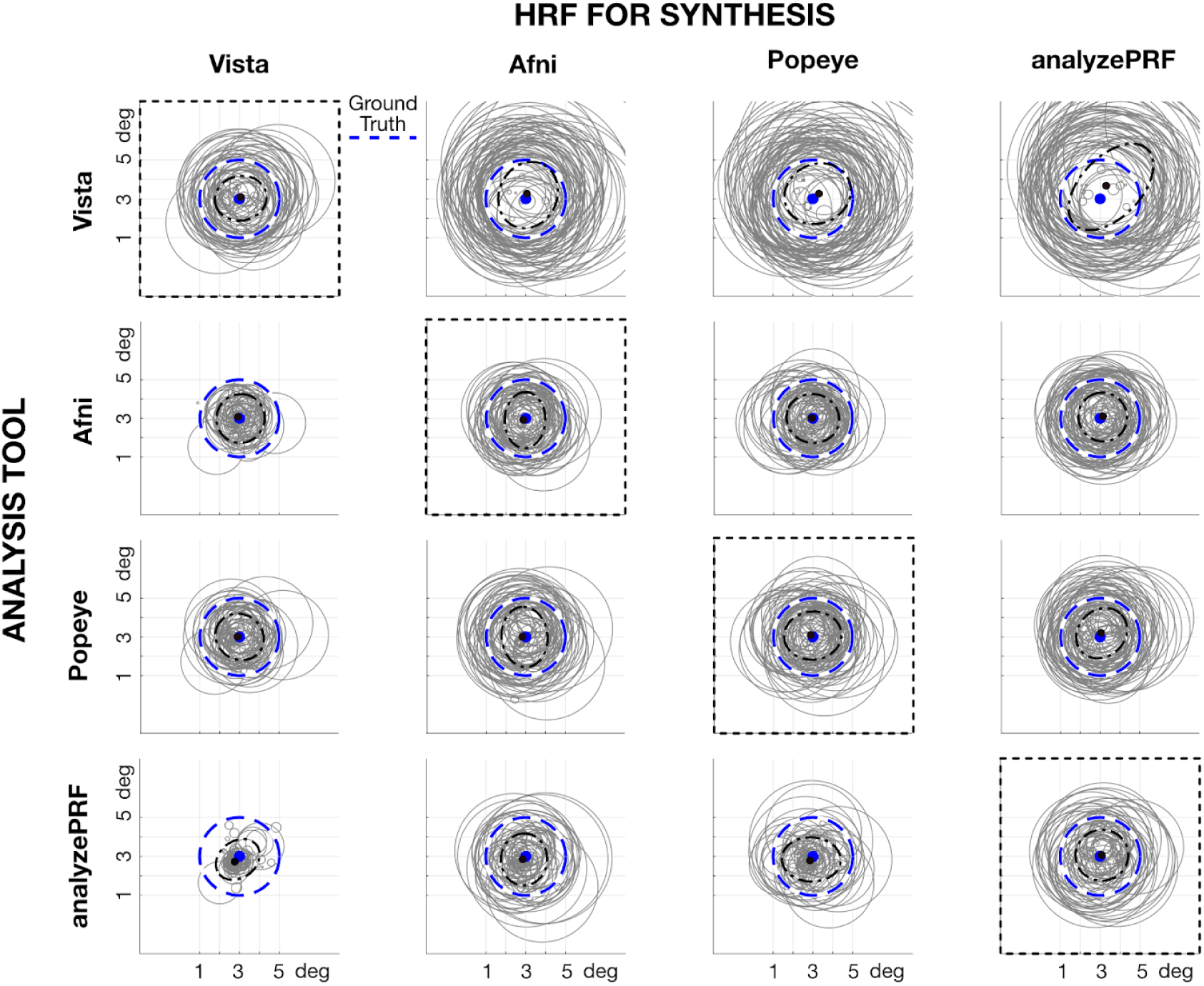
Noise analysis and HRF dependence. Each column shows the HRF used in pRF-Synthesize to simulate the BOLD time series. The prf-Synthesize tool created BOLD signals with the typical noise level (mid level, see Figure 4) and a circular RF centered at (3,3) deg and 2 deg radius (dashed blue circle). Each row corresponds to the pRF-Analyze tool, with its default HRF, that was used to analyze the data. The gray circles show the central 90% of the estimated RFs. The central ellipsoid includes 90% of the estimated centers (dashed-dot, black). The central black dot shows the median center location. The HRF used in the synthesis matches the HRF assumed in the analysis in the plots along the diagonal (dashed black rectangles). Above the diagonal, the synthesis HRF is narrower than the analysis HRF; below the diagonal the opposite is true.

In fMRI experiments we have no control over HRFs, which differ between people and across cortical locations. We created a simulation using a range of HRFs for each tool. The confidence interval of the pRF radius is almost ±2 degrees. This value can change depending on the size of the ground truth pRF size, but the general pattern is the same.

## Discussion

The validation framework exposed meaningful implementation differences. The differences that we discovered would lead investigators to report different quantitative parameter estimates from otherwise equivalent data. This finding illustrates the value of the validation framework in making meaningful comparisons across labs. We can reduce conflicts that arise when two labs report differences that arise from the prf-Analyze tool rather than empirical measurements.

We illustrate the framework with population receptive field (pRF) tools because there are many: We found 11 public pRF implementations (https://github.com/vistalab/PRFmodel/wiki/Public-pRF-implementations). It is likely that extending the validation framework to test the additional prf-Analyze tools will reveal other meaningful differences. We hope that making the ground truth datasets and the synthesis and analysis containers available to the community will help software developers who are creating new methods, and researchers to have confidence in their tool choice.

More work remains to be done for the pRF-Validation, but we can already report two meaningful findings. First, we discovered and reported issues in some implementations. Second, there is a significant dependence of the pRF size estimate on the HRF, and the HRF model differs significantly between implementations. This should be accounted for when comparing numerical estimates of pRF size using the different tools. There is no significant effect on position estimates of the pRF. These findings illustrate the value of using a software validation frameworks.

### HRF dependence

We found that the default HRF assumed in the analysis tool can have a large effect on the size of the estimated pRF. The effect on eccentricity and polar angle was negligible. This is due in part to the stimulus design, which included bar sweeps in both directions along several axes (e.g., left-to-right and right-to-left). If the stimulus were not symmetric (e.g., containing left-to-right sweeps but not right-to-left), then a mismatch between the HRF assumed by the analysis tool and synthesis tool would have resulted in larger errors in the estimated pRF center position.

The HRF impact on pRF size estimates raises two main concerns. The most obvious one is that we cannot compare the absolute size results across tools with different HRF assumptions; the comparison only makes sense within the same analysis tool. The second and more concerning is that even with the same tool, two participants may have different HRFs, or the same person may have different HRFs in different locations in the brain [22]. This HRF dependence has been noted by other investigators who have adopted various means to estimate individual participant HRFs [23].

There may be many approaches to mitigating the HRF dependence. We describe approaches here.

#### Computational mitigation

One approach to reducing the impact of the HRF is to estimate it. There are many ways to approach the HRF estimation. Some investigators allow every temporal sample to be a free parameter [24,25]. Measuring a subject’s HRF in a separate experiment adopts this approach. Other investigators parameterize the range of HRF possibilities, using a model with a few parameters (see [22] for a review on alternatives). Some investigators use independent experiments to estimate the HRF and the pRF; others perform a joint estimation. These alternatives recognize the possibility that the HRF is stimulus dependent, in which case the estimate from an impulse may not generalize to the estimate from the retinotopic data.

Accurately estimating the HRF in every voxel is challenging and vulnerable to noise. Two of the prf-Analyze tools, mrVista and Popeye, have an option to fit the HRF parameters in addition to the pRF parameters. Preliminary validation tests suggest that the effectiveness of the HRF fitting option, although positive, was small. This is a good direction for future exploration.

#### Empirical mitigation

A second approach to reducing the impact of the HRF is to change the stimulus. Slowing the bar sweep across the screen reduces the impact of the HRF on the estimated pRF size (Figure 9). The three rates shown in the figure are representative of values found in recently published measurements [26–28]. When the bar sweep is slowest (38 sec), the HRF mismatch has much less effect on the estimated pRF size. Selecting this approach substantially increases the amount of time required to obtain the measurements.

**Figure 8.**
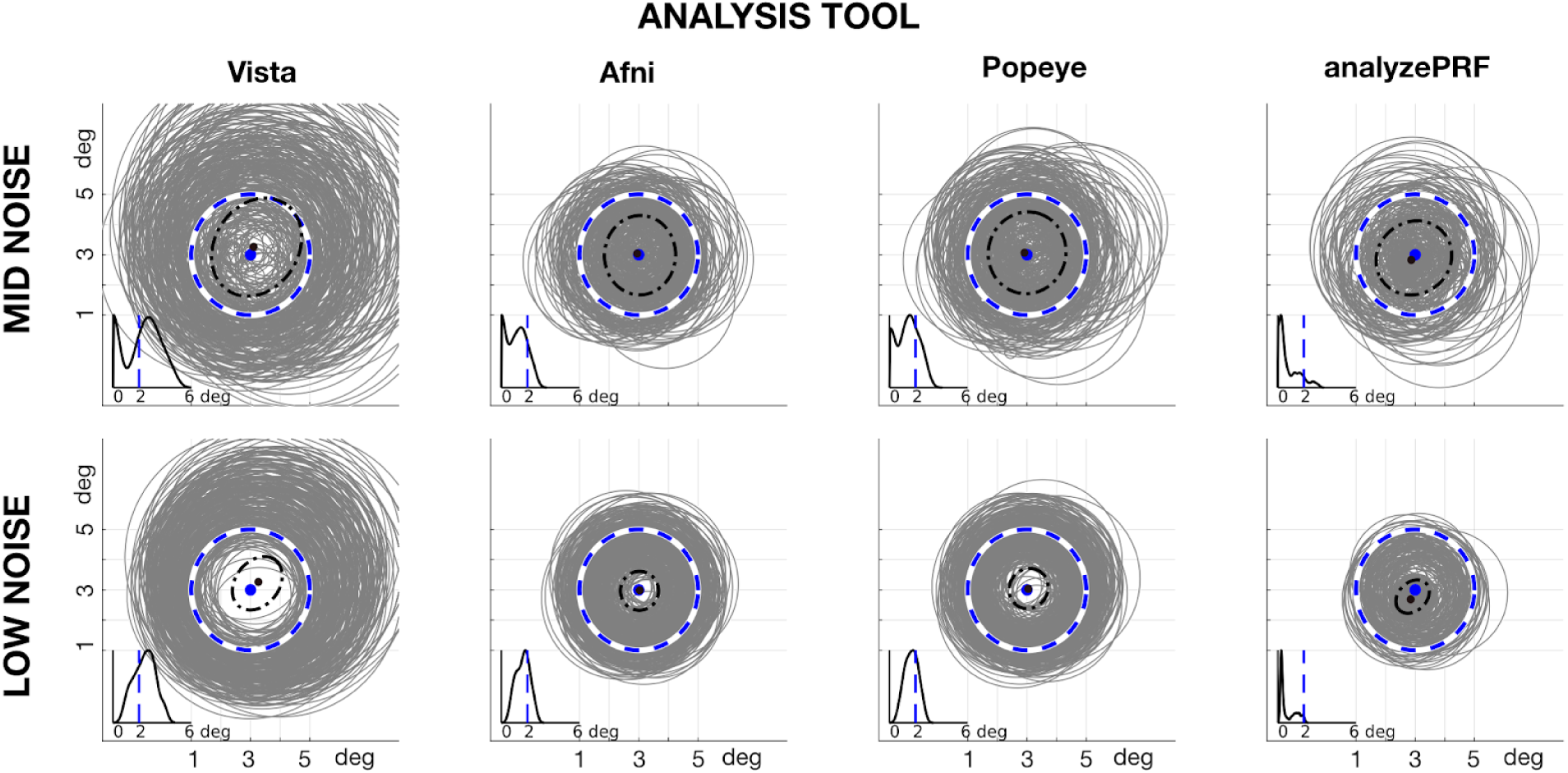
HRF variation. The synthesized circular pRF was centered at (3,3) deg with a radius of 2 deg. The data were simulated with prf-Synthesize 400 times, and each time using one of the four prf-Analyze HRFs to synthesize the data. The estimated circular pRFs (gray circles) were calculated using the default HRF implemented in each tool. Outliers are eliminated by showing only fits within the 90th percentile of the pRF size estimates. The inset shows the probability distribution of the pRF sizes, where the blue vertical line shows the ground truth (2 deg).

**Figure 9.**
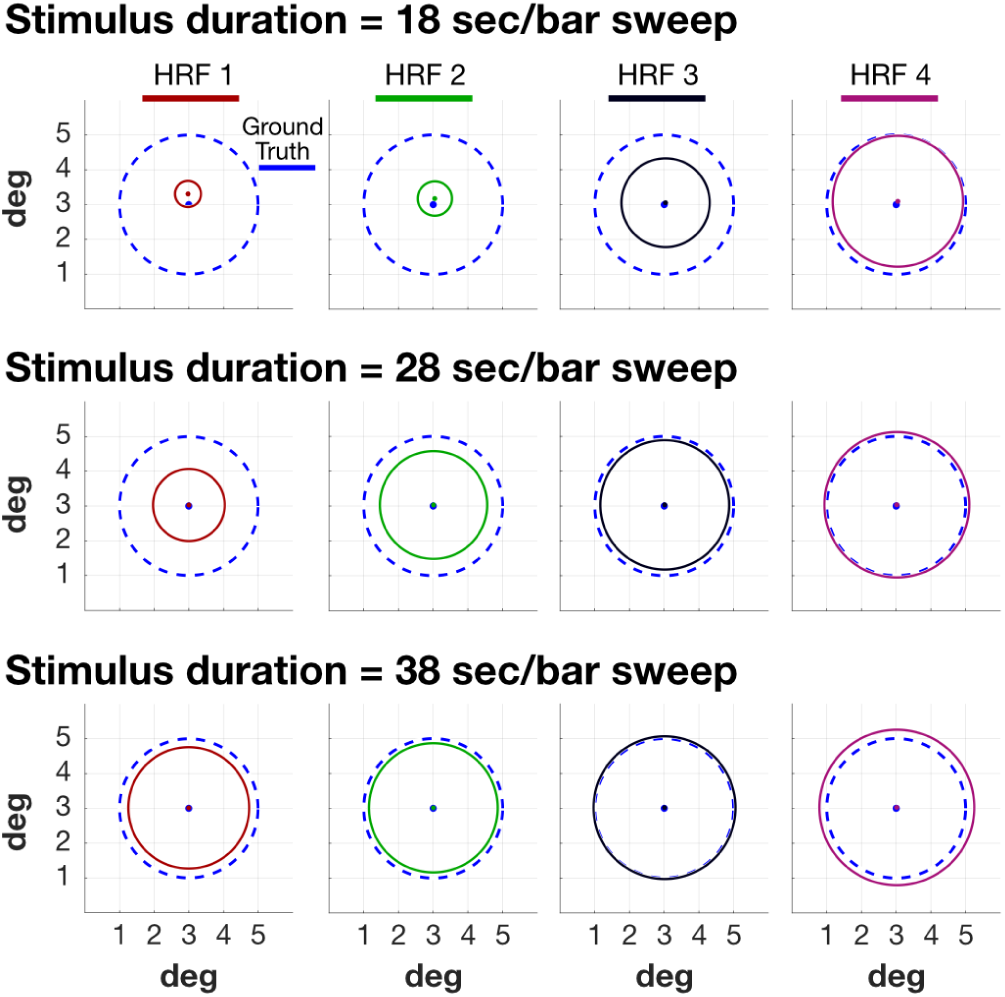
Stimulus duration reduces the impact of HRF variation. Slowing the bar movement through the visual field reduces the HRF mis-match impact. The three rows extend the calculation in Figure 6, using three bar speeds: 18, 28 and 38 sec per sweep. Other details as in Figure 6.

Randomizing the position of the bar, rather than sweeping it across the visual field, also reduces the impact of the HRF on the estimated pRF size (Figure 10A). The benefit of position-randomization depends on the specific randomization. It may be possible to find a specific randomized pattern that optimizes the validity of the pRF size estimate for a particular HRF and pRF size.

**Figure 10.**
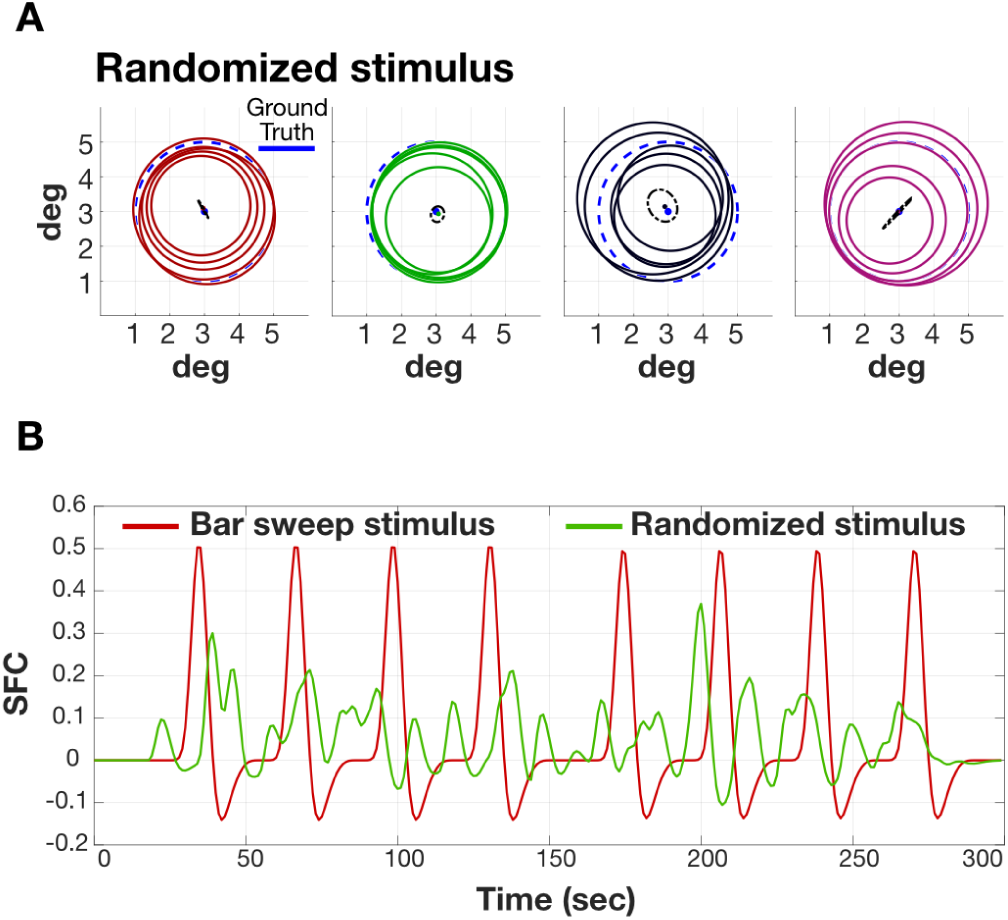
Stimulus randomization reduces the impact of HRF variation. (A) Randomizing the bar positions in time reduces the HRF mis-match impact. This figure extends the calculation in Figure 6 and Figure 9, using 5 different randomization seeds. Other details as in Figure 6. (B) Noise-free time series corresponding to bar-sweep stimulus (red) and randomized stimulus (green). The SNR will be lower due to the randomization of the stimuli.

Choosing a randomized bar position, rather than a sweep pattern, comes at the cost of a significant loss of contrast in the BOLD response (lower signal-to-noise, Figure 10B). For the sweep patterns the bar systematically passes through the pRF of each voxel, driving the response to its maximum. Randomizing the position reduces the amount of time that the bar is positioned over the pRF and consequently reduces the response contrast. The loss of response contrast increases the amount of time required to obtain estimates with a particular confidence interval.

### Outliers

Because prf models are not linear with respect to their parameters, the solutions are found by non-linear search algorithms. The algorithms sometimes return non-optimal local minima. It is not unusual for many of the algorithms we tested to return extreme cases that are outliers. Some of the algorithms mitigate this problem by placing a bound on the returned estimate. Other algorithms incorporate resampling, running the search with different starting points and testing for agreement between the returned parameters.

The simulations included 100 repetitions for the same voxel, with randomized noise. In these simulations the median result is close to the real solution. We assume that this approximates what is observed with real data; the median of the results yield the right solution but that there are outliers. In practice we suggest that repeating the pRF analysis with different seeds three or four times will approximate the median solution. Alternatively, it is possible to run the analysis by adding a small amount of noise to every real fMRI time series and report the median result.

There is another approach which we have not yet seen implemented. In many parts of visual cortex nearby voxels will have similar pRF parameters. When this is known to be true, it is possible to increase the effective SNR of the measurements by including an expectation that the estimates from nearby voxels will be similar.

### Guiding experimental design

We are using the tool as a simulator to design new types of stimuli and experiments prior to data acquisition. For example, here we used the tool as a simulator to identify the two empirical methods for mitigating the unwanted impact of the unknown HRF. This approach may also be useful when making a stimulus and experimental design choices. The prf-Synthesize and prf-Analyze tools are helpful for assessing whether the design will support a meaningful test of a specific hypothesis.

### Limitations

Validation testing provides an opportunity to consider several limitations of the population receptive field methods. We comment on these first, and then we make some observations about the validation framework itself.

#### Noise and HRF stimulus-dependence

We estimated the noise in the BOLD signal from an on/off experiment. We are not sure that this noise estimate is perfectly matched to the noise in the retinotopic measurement. This is true for both the signal amplitude and the correlation. In particular, the time series are synthesized independently. We expect that adjacent voxels in the BOLD measurements generally have correlated noise. Incorporating this spatial correlation may be useful. Moreover, we assumed additive noise but there may also be multiplicative noise, perhaps due to variations in attention or arousal [29]. In the future, it may be useful to express certain types of noise with respect to the input units, such as eye movements, rather than output units (% BOLD). As noted earlier, the HRF may be stimulus dependent, it may vary across visual cortex, and it may differ significantly between participants.

#### prf-Analyze parameter selection

We made a best-faith effort to optimize the configuration parameters, sometimes consulting with the authors. Further, in creating the Docker containers we use a JSON file that enables the user to configure the prf-Analyze tools. There is room to improve the documentation of the parameters in order to help researchers use the tools more effectively. The validation framework supports the documentation because users can vary parameters and explore the consequences.

#### Creating containers

Exploring the validation framework is simple for most users: they need only a text editor and Docker. It would be desirable to make contributions to the validation framework equally simple. At present contributing a new prf-Analyze tool involves the following steps. If the pRF tool is implemented in Python, the developer must configure a Docker container that includes all of the required dependencies. If the tool is implemented in Matlab, the source code must be compiled and placed in a Docker container that includes the Matlab run-time environment (we provide this for Matlab 2018b). In both cases, the tool must read inputs and write outputs according to the BIDS standard. We provide functions for this purpose (see the prf-Validation Guide).

## Conclusion

Scientists recognize the tension between using established methods and extending these methods to make new discoveries. Scientists must have confidence in the validity of the methods which advocates for a conservative approach to software use: select tools that have been widely tested within the community. But science is also about exploring new ideas and methods which advocates for a more liberal approach: try new methods that have the potential to expand our understanding. In practice, most scientists are advocates of both creativity and caution.

It is this tension that motivates us to work with the validation framework described in this paper. We find it too confining to follow the oft-repeated suggestion that investigators rely upon a few agreed upon packages. But if we are to explore, it is essential to use a testing framework that can help (a) developers to test new software, (b) research scientists to verify the software’s accuracy and compare different implementations, and (c) reviewers to evaluate the methods used in publications and grants. A public validation framework, using reproducible methods and shared ground-truth dataset, is a useful compromise between caution and creativity. In the future we plan to contribute more ground truth datasets and analyses of pRF tools, and we also hope to add validations of additional neuroimaging tools. We hope to work with others to extend the validation framework to validate tools in diffusion-weighted imaging, quantitative MRI, resting-state, and anatomical methods.

## Acknowledgements

Supported by a Marie Sklodowska-Curie (H2020-MSCA-IF-2017-795807-ReCiModel) grant to G.L.-U. and NIH grants supporting N.C.B. and J.W. (EY027401, EY027964, MH111417).

## Appendix

### prf-Validation guide

The modular architecture of the testing framework is based on containers and the Brain Imaging Data Structure (BIDS) organization. Consequently means the tool runs on most platforms without compilation (Figure Appendix 1). The synthesize-analyze-report pipeline is executed with a few command lines. The parameters that control the pipeline execution are specified by editing the three configuration files. More detailed instructions for installing the framework are maintained on a website (https://github.com/vistalab/PRFmodel/wiki).

**Appendix Figure 1.**
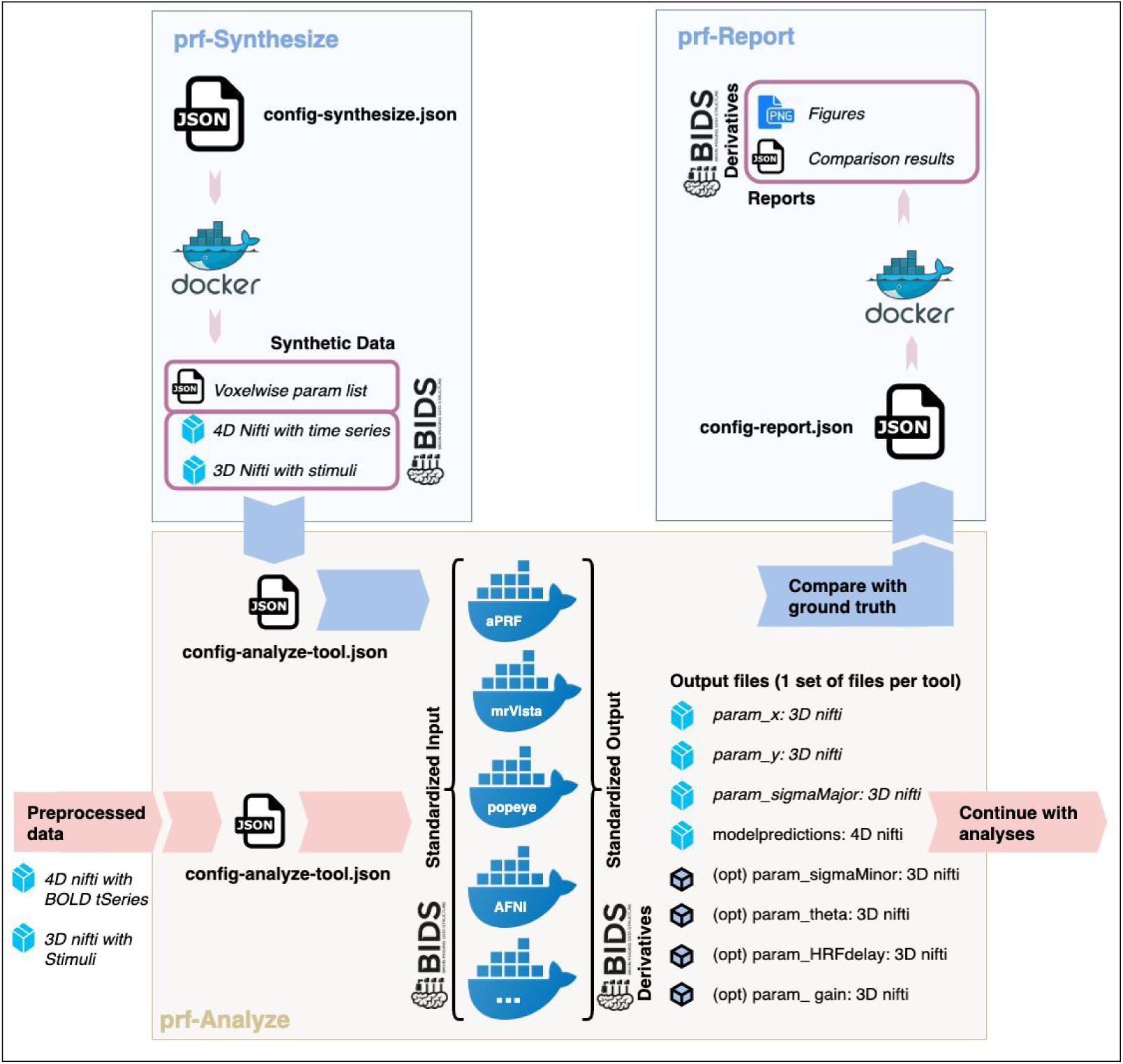
Detailed working steps of the 3 element validation framework. *prf-Synthesize:* using a default json config file, allows the generation of a BIDS folder structure with the synthetic data and the parameters of every time series. *prf-Analyze:* Takes synthetic or real fMRI datasets in BIDS format, adapts them to different implementations of tools, and generates a BIDS derivatives output with uniformized output parameters. *prf-Report:* takes the synthetic dataset output from prf-Analyze and the voxelwise parameter definition with the ground truth from prf-Synthesize, and calculates goodness of fit reports.

#### Installation

Docker must be installed in your system. The tools are installed with “docker pull” command lines. See installation instructions in the Github wiki (https://github.com/vistalab/PRFmodel/wiki#how-to-install).

#### prf-Synthesize

The prf-Synthesize container is invoked by the command ‘run_prfsynth’ or with a direct docker run call. That command takes two arguments: (a) the name of the configuration file (json format), and (b) a string with the full path to the output directory. The output file names are assigned by the container. A default configuration file can be created by running the container with the arguments ‘empty’ and the path to the output folder. In that case the container will return a json called `‘prfsynth-config-defaults.json’ in the $PWD folder`. Using a text editor, the user modifies the json file, specifying the parameters for the synthetic BOLD time series. Detailed instructions on how to edit the file can be found in the GitHub wiki (https://github.com/vistalab/PRFmodel/wiki/prf-Synthesize:-how-to-edit-json-file). After the json file has been edited, the user runs the Docker container again to create the synthetic data in the output folder.

The container is designed to create a grid of all combinations of the configuration parameters. For example, if the configuration file includes two center locations for x and y, say 0 and 5, the container generates 4 synthetic BOLD series with centers at [0,0], [0,5], [5,0] and [5,0]. If two HRFs are specified in the configuration file, say ‘popeye_twogammas’ and ‘afni_spm’, the number of synthetic BOLD time series doubles to 8; the four center locations times two HRFs. If we the configuration parameter ‘repeats’ is set to 100, 100 noisy samples are generated for every parameter combination. The documentation in the GitHub wiki contains a more detailed description.

Upon successful completion the container creates a new folder called ‘BIDS’ in the output directory. The files are organized following the BIDS guidelines. A NIfTI file with the BOLD time series is in the *subject/session* directory, a NIfTI file defines the stimulus in the *Stimulus* directory, and a json file defining parameters used to generate each BOLD time series is in the *derivatives/prfsynth/subject/session* directory.

The files written out by prf-Synthesize are used by the analyze and report elements of the framework.

#### prf-Analyze

The prf-Analyze container is invoked by the command ‘run_prfanalyze-METHOD’. That command takes two arguments: (a) the name of the configuration file (json format), and (b) a string with the full path to the base folder of the BIDS directory. For example, for the ‘analyze prf’ model the command is ‘run_prfanalyze_aprf’. The output file names are assigned by the container. We have implemented multiple methods, and we hope that more will be contributed.

To initialize the configuration file for a prf-Analyze method, the user runs the Docker container with the `‘empty’` parameter and the full path to the base folder of the BIDS directory. The Docker container initializes a json configuration file; in this example the file name will be `‘prfanalyze-configuration-defaults.json’`. The parameters in the prf-Analyze configuration file are tool dependent. The parameters in the file need to be set, including the data directory and the required parameters to run the pRF analysis, see here for detailed instructions (https://github.com/vistalab/PRFmodel/wiki/prf-Analyze:-how-to-edit-json-file). Once the json file has been edited, we call the Docker container again, providing the full path to the json file and the pull path to the output folder.

Upon successful completion of the prf-Analyze container, a new folder with the tool name is created in theBIDS *derivatives* directory (e.g. *derivatives/prfanalyze-afni)*. This folder contains the *subject/session* folder with a set of NIfTI files, one per each of the parameters estimated by the tool. Some parameters are mandatory (see Figure 1):

1. sub-subject_ses-session_task-prf_x0.nii.gz: NIfTI file containing a predicted x parameter per voxel.
2. sub-subject_ses-session_task-prf_y0.nii.gz: NIfTI file containing a predicted y parameter per voxel.
3. sub-subject_ses-session_task-prf_sigmamajor.nii.gz: NIfTI file containing the radius of the circular RF, in the case of elliptical fitting, it will be the sigmaMajor.
4. sub-subject_ses-session_task-prf_sigmamajor.nii.gz: NIfTI file containing the radius of the RF.
5. sub-subject_ses-session_task-prf_modelpred.nii.gz: NIfTI file containing the BOLD time series prediction from the tool. It should be scaled back at the input BOLD series to allow direct comparisons (variance explained, for example).

Other METHOD-specific NIfTI files will be created. For example, if the simple circular AFNI implementation is used, an additional NIfTI file specifies the gain. If the elliptical AFNI implementation is used there are additional files for the gain, sigmaMinor file and Theta angle value that expresses the inclination of the sigmaMajor axis with respect to the horizontal axis.

These files are ready to be used in the prf-Report part of the validation framework.

#### prf-Report

The usage is analogous to the other blocks. We run it with the ‘empty’ parameter to obtain the default config-report json file, named `‘prfreport-configuration-defaults.json’`. Then we run it with the edited config file to perform the analysis. See detailed instructions (https://github.com/vistalab/PRFmodel/wiki/prf-Report:-how-to-edit-json-file) about editing the config file. We place the analyses in the BIDS derivatives folder, with a name such as *derivatives/prfreport/subject/session*.

##### Visualization and customization of the main elements

The Docker containers were implemented to simplify running the code on most platforms. If one desires to examine the components of the calculations one-by-one and check individual results or hypotheses, it is also possible to run the code manually within the Matlab command line interface. Download and set-up the code repository (http://github.com/vistalab/prfmodel) and follow the online instructions. Check the guide in the wiki for further detail (https://github.com/vistalab/PRFmodel/wiki/Visualization-and-customization-of-the-main-elements-in-Matlab).

##### Public pRF implementations

We maintain a list of public pRF analysis tool implementations here (https://github.com/vistalab/PRFmodel/wiki/Public-pRF-implementations).

